# CiliAI: Automated segmentation and compartment-specific fluorescence quantification of primary cilia in confocal microscopy images

**DOI:** 10.64898/2026.06.05.727888

**Authors:** Eliza Karapetian, Christoph Gerhardt, Emrah Nazif, Thorsten Pfirrmann

**Affiliations:** Health and Medical University, Potsdam, Germany; Martin-Luther University, Halle (Saale), Germany; Datamarkin, Marseille, France

**Author notes:** These authors contributed equally to this work (Eliza Karapetian, Christoph Gerhardt).

**Keywords:** Vision AI, Machine Learning, Artificial Intelligence, primary cilium, ciliopathies, image analysis, cancer diagnosis

## Abstract

Primary cilia regulate essential signaling pathways controlling cell proliferation, differentiation, and tissue homeostasis. Quantitative analysis of ciliary morphology and compartment-specific protein localization by confocal microscopy is labor-intensive, user-dependent, and difficult to scale, particularly for multiplexed 3D image datasets. Here, we present CiliAI, a web-based deep-learning workflow for automated detection, substructure segmentation, and quantitative analysis of primary cilia in confocal microscopy images. CiliAI identifies ciliary substructures including the basal body, transition zone, and axoneme from multiplexed 3D image stacks and performs automated measurements of cilium length and compartment-specific fluorescence intensity. In NIH-3T3 cells, automated cilium length measurements showed close agreement with manual quantification and no statistically significant difference between methods (mean difference −0.214 µm, p = 0.213). Automated fluorescence analysis reproduced previously reported reductions in transition zone-associated Cep290 signal intensity in Rpgrip1l-deficient cells and identified the absence of significant Rpgrip1l accumulation changes in Rmnd5a-deficient cells. Automated processing reduced analysis time from days of manual quantification to minutes. Together, these findings establish CiliAI as an automated framework for quantitative analysis of ciliary morphology and compartment-specific protein abundance in confocal microscopy datasets.

## 1 Introduction

Primary cilia are conserved sensory organelles present on most vertebrate cells that coordinate signaling pathways controlling proliferation, differentiation, polarity, and tissue homeostasis ^1-5^ Structurally, the primary cilium consists of distinct subcompartments including the basal body (BB), transition zone (TZ), and axoneme, which regulate ciliary assembly, signaling, and protein trafficking ^6,7^.

Defects in primary cilia cause a heterogeneous group of disorders collectively termed ciliopathies ^8-13^. Classical ciliopathies are characterized by heterogeneous clinical manifestations including kidney disease, retinal degeneration, and skeletal abnormalities ^3,14^. Beyond classical ciliopathies, altered ciliary signaling and protein localization have also been implicated in cancer and neurodegenerative diseases ^15-19^. Protein abundance within ciliary subcompartments is regulated by trafficking and proteostasis mechanisms, and disruption of these processes contributes to ciliopathy-associated phenotypes ^20-22^. These findings highlight the need for reproducible methods capable of measuring compartment-specific protein abundance within primary cilia subcompartments.

Quantitative analysis of ciliary morphology and compartment-specific fluorescence signals from multiplexed confocal z-stacks remains labor-intensive and user-dependent, requiring manual region-of-interest selection across multiple image planes ^23-25^. Automated analysis of primary cilia in confocal microscopy datasets is complicated by the small size and morphological variability of ciliary structures, heterogeneous fluorescence intensities, and the requirement for subcompartment-specific segmentation across multiple imaging channels.

Previous tools including CiliaQ ^26^ and ACDC ^27^ enabled automated cilia detection and morphometric analysis, while newer workflows support automated ciliary subdomain analysis ^28^. However, automated integration of cilia detection, substructure segmentation, and compartment-specific fluorescence quantification from multiplexed 3D confocal datasets remains limited.

Here, we present CiliAI, a web-based workflow for automated detection, substructure segmentation, and compartment-specific fluorescence quantification of primary cilia in confocal microscopy datasets. CiliAI processes 3D image stacks to identify ciliary substructures including the basal body, transition zone, and axoneme, and generates quantitative measurements of cilium length and fluorescence intensity. By standardizing compartment-specific analysis of ciliary morphology and fluorescence intensity, CiliAI provides a reproducible workflow for quantitative analysis of primary cilia in confocal microscopy datasets.

## 2 Results

### Automated workflow for cilia detection and substructure segmentation

**Figure 1A** illustrates the major structural subcompartments of the primary cilium, including the basal body (blue), transition zone (red), axoneme (green), and ciliary membrane (orange), shown schematically and by representative confocal microscopy. **In Figure 1B**, the basal body was labeled using anti-γ-tubulin antibodies (blue), the transition zone using anti-RPGRIP1L antibodies (red), and the axoneme using anti-detyrosinated tubulin antibodies (green).

**Figure 1:**
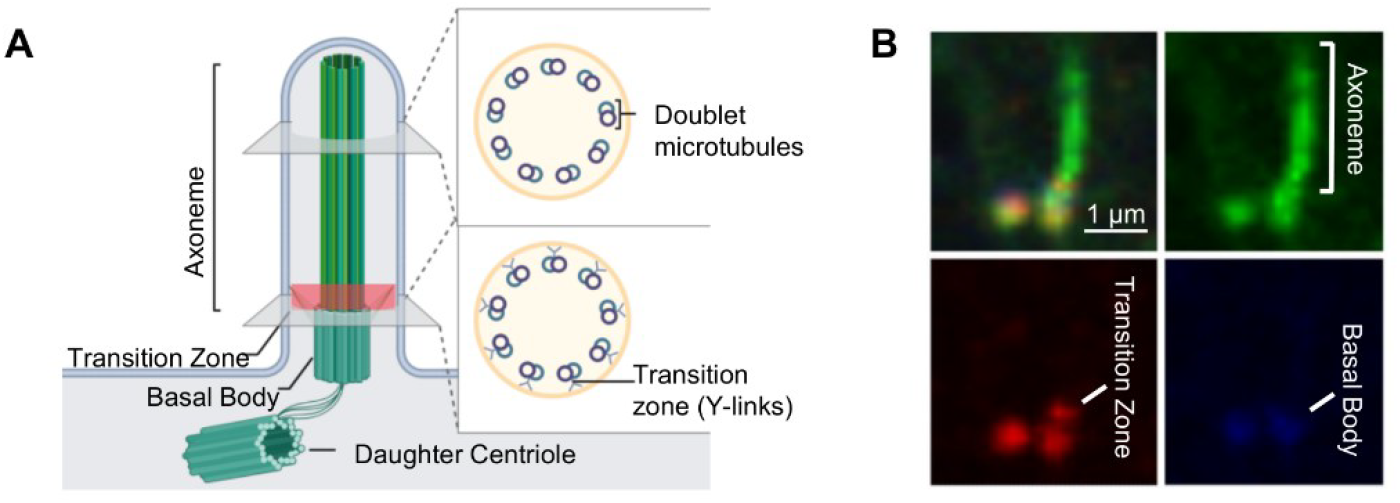
Structure and subcompartments of a primary cilium. (A) Schematic representation of a primary cilium showing the axoneme, transition zone, basal body, and daughter centriole. Insets illustrate the arrangement of microtubule doublets within the axoneme and Y-link structures of the transition zone. (B) Representative confocal microscopy images highlighting the same substructures: axoneme (green, anti-acetylated tubulin), transition zone (red, anti-Rpgrip1l), and basal body (blue, anti-γ-tubulin). Scale bar: 1 µm.

The CiliAI workflow for automated primary cilia analysis is illustrated schematically in **Figure 2A**. Confocal microscopy datasets are processed through sequential stages comprising cilia detection, substructure segmentation, and compartment-specific fluorescence quantification. Detected ciliary substructures such as the basal body, transition zone, and axoneme are identified within processed image stacks. In the final step, cilia-specific parameters including cilium length and compartment-specific fluorescence intensity are quantified across multiple fluorescence channels. To improve detection of small ciliary structures, confocal images were subdivided into overlapping regions of interest followed by higher-resolution analysis of detected cilia (**Figure 2B**).

**Figure 2:**
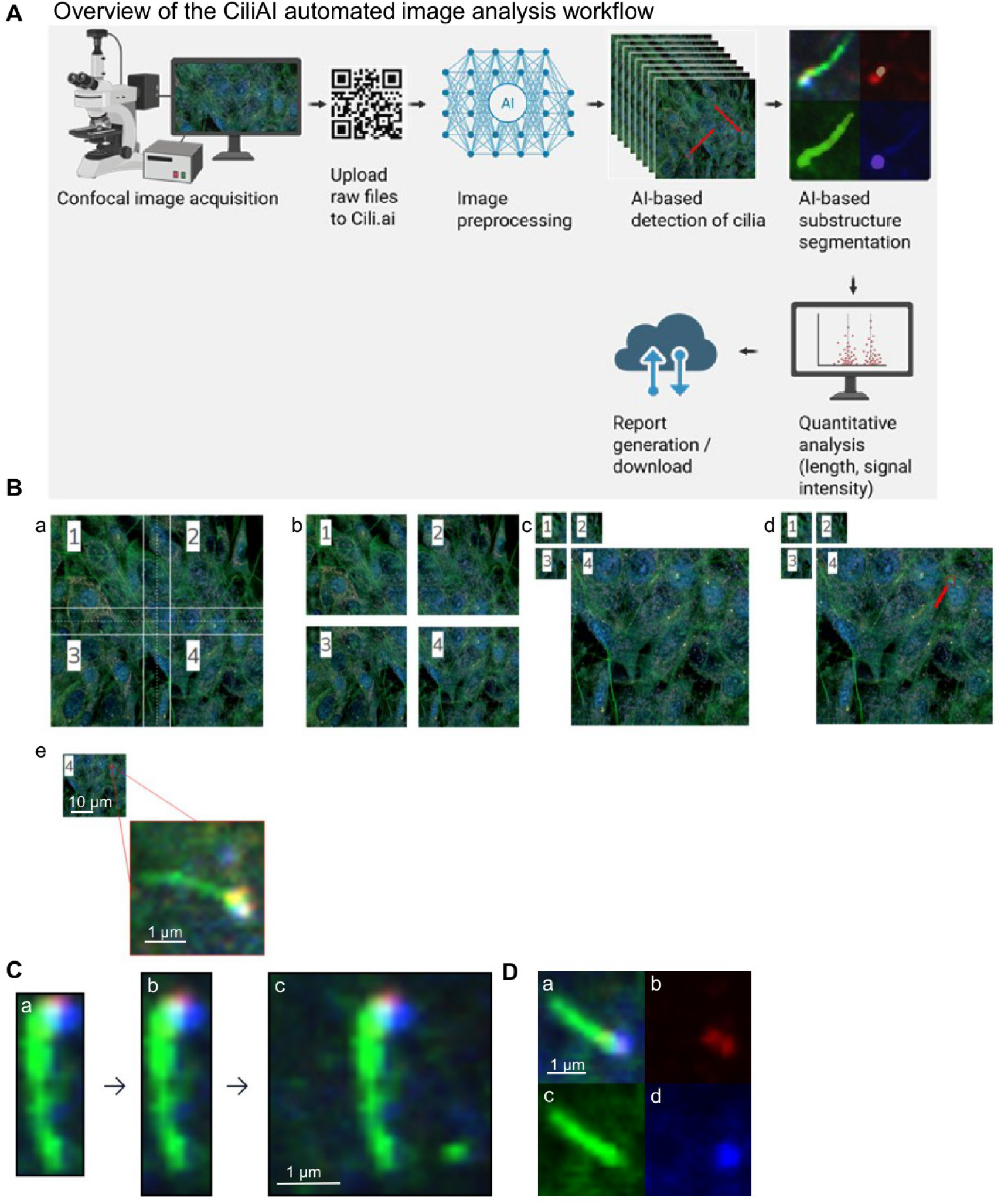
Automated image analysis workflow in CiliAI. (A) Overview of the CiliAI pipeline from confocal image acquisition and file upload to AI-based preprocessing, cilia detection, and substructure segmentation, till quantitative analysis and automated report generation. (B) Image tiling and region-of-interest segmentation: each confocal image is divided into quadrants to ensure accurate detection across z-stacks; quadrant 4 highlights a detected cilium (red arrow). (C) Cropping and upscaling workflow showing progressive magnification and resolution enhancement for substructure analysis. (D) Segmentation of ciliary substructures visualized in composite (RGB) and single-channel views: transition zone (red), axoneme (green), basal body (blue). Scale bars: 1 µm (B), 0.5 µm (C–D). AI = artificial intelligence; ROI = region of interest.

CiliAI generates merged detection maps across confocal z-stacks and assigns confidence scores to detected cilia (**Figure 3A**). For each detected cilium, the workflow outputs quantitative parameters including cilium length, fluorescence intensity, and segmented substructure areas (**Figure 3B**). Quantitative outputs are exported in tabular format for downstream analysis (**Figure 3B**). Segmented ciliary substructures are visualized as composite and channel-specific overlays (**Figure 3C**).

**Figure 3:**
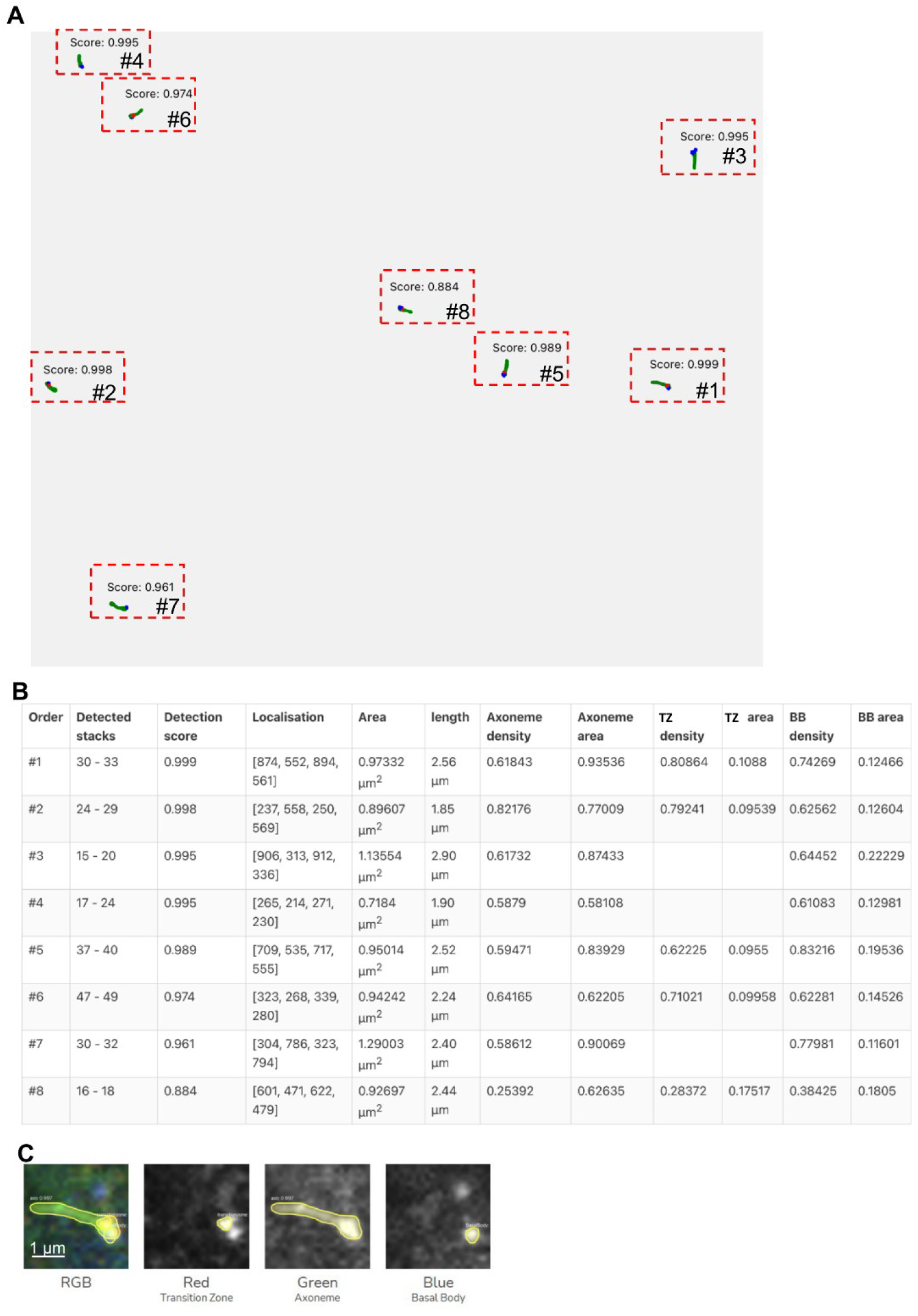
Automated detection, quantification, and visualization of primary cilia using CiliAI. (A) Representative top-view 2D detection map showing identified primary cilia (outlined in red) and corresponding detection scores. Each cilium is labeled with its detection order. Detections with confidence ≥ 0.5 are considered true positives. (B) Quantitative output table summarizing per-cilia parameters including detection score, cilium length, and intensity and area measurements for the axoneme, transition zone (TZ), and basal body (BB). (C) Example of substructure segmentation displayed in composite (RGB) and individual channels: red = transition zone, green = axoneme, blue = basal body. TZ = transition zone; BB = basal body; confidence scores > 0.5 = true detections.

### AI model performance for cilia detection and substructure segmentation

Both models were evaluated on a held-out validation set, split at the level of source 3D image stacks so that no tile or cilium from a validation stack appeared in training (Section 4.7). During training, total loss decreased and validation performance stabilized for both the detection and segmentation models (**Figure 4A,C**).

**Figure 4.**
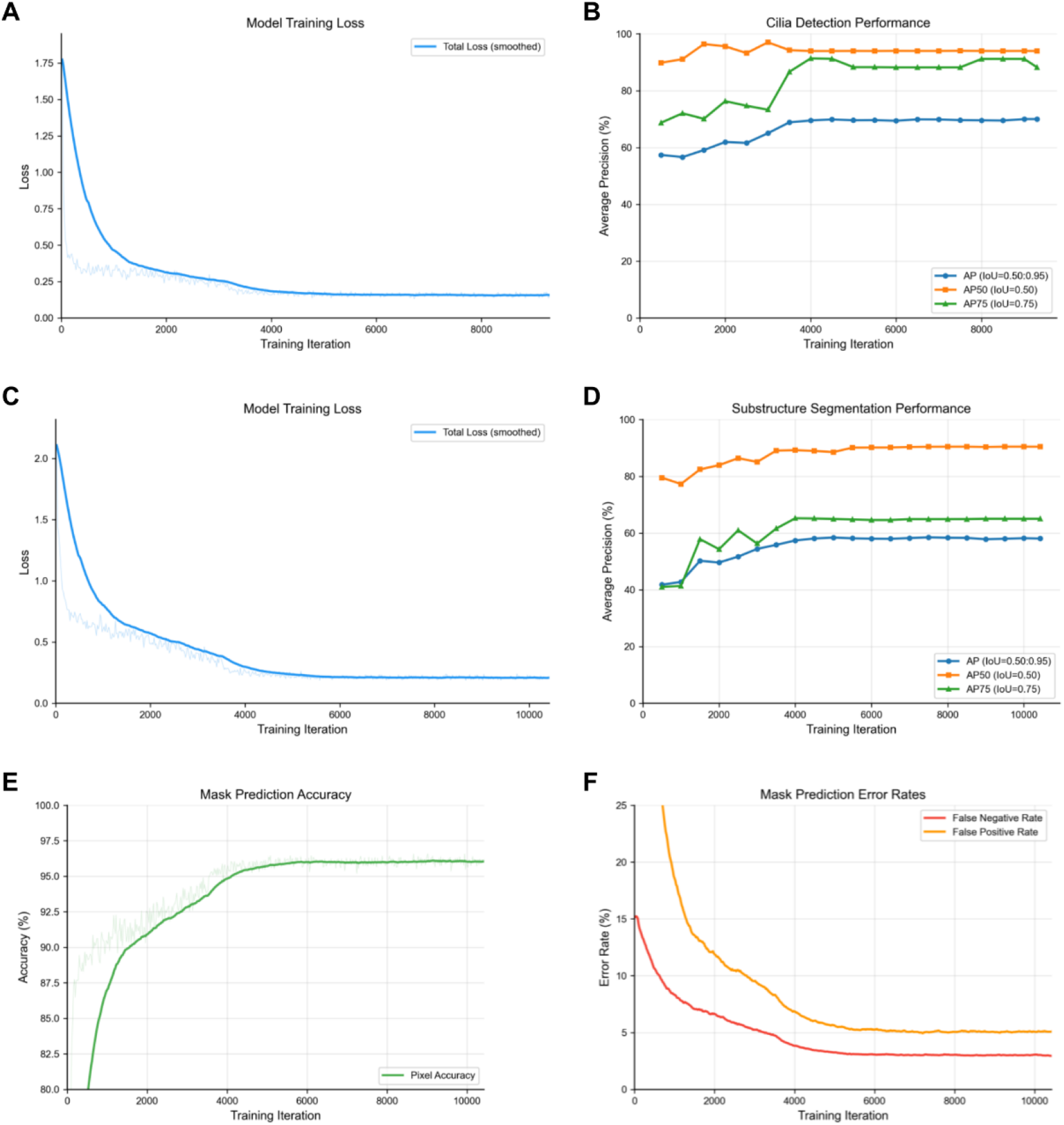
AI model performance during cilia detection and substructure segmentation. (A) Training loss curve for the cilia detection model. (B) Detection performance measured as AP50, AP75, and mAP50–95 during training. (C) Training loss curve for the substructure segmentation model. (D) Segmentation performance measured as AP50, AP75, and mAP50–95 during training. (E) Pixel-level mask prediction accuracy during training. (F) False-positive and false-negative mask prediction error rates during training.

For cilia localization, the detection model achieved a segmentation AP50 of 97.0% and a bounding-box AP50 of 94.0%, while the mean average precision across IoU thresholds from 0.50 to 0.95 (AP) reached 67.5% (segmentation) and 70.0% (bounding box) (Figure 4B). The lower AP relative to AP50 is consistent with the difficulty of exact boundary localization for small, morphologically variable structures, where object-level detection is more reliable than pixel-precise delineation.

For substructure segmentation, the model reached a segmentation AP50 of 90.5% and a mean AP of 58.1% across the three ciliary compartments (Figure 4D). Per-class segmentation AP was highest for the axoneme (74.8%) and basal body (65.4%), and lower for the transition zone (34.0%), consistent with the transition zone being the smallest and most morphologically variable compartment and therefore the most challenging to segment precisely. Together, these results indicate that both models achieved performance sufficient to support downstream morphometric and compartment-specific fluorescence quantification, while identifying transition-zone segmentation as the principal target for future refinement.

### Validation of automated morphometric and compartment-specific fluorescence quantification

To evaluate the performance of CiliAI, automated measurements were compared with conventional manual analysis methods. Cilium length measurements were first evaluated in NIH-3T3 cells. To measure the length of cilia, we fixed and stained WT NIH-3T3 cells using anti-detyrosinated alpha-tubulin (axoneme), anti-γ-tubulin (basal body) and anti-RPGRIP1L (transition zone). Automated cilium length measurements showed close agreement with manual quantification and no statistically significant difference between methods (mean difference −0.214 µm, p = 0.213; **Figure 5A**).

**Figure 5:**
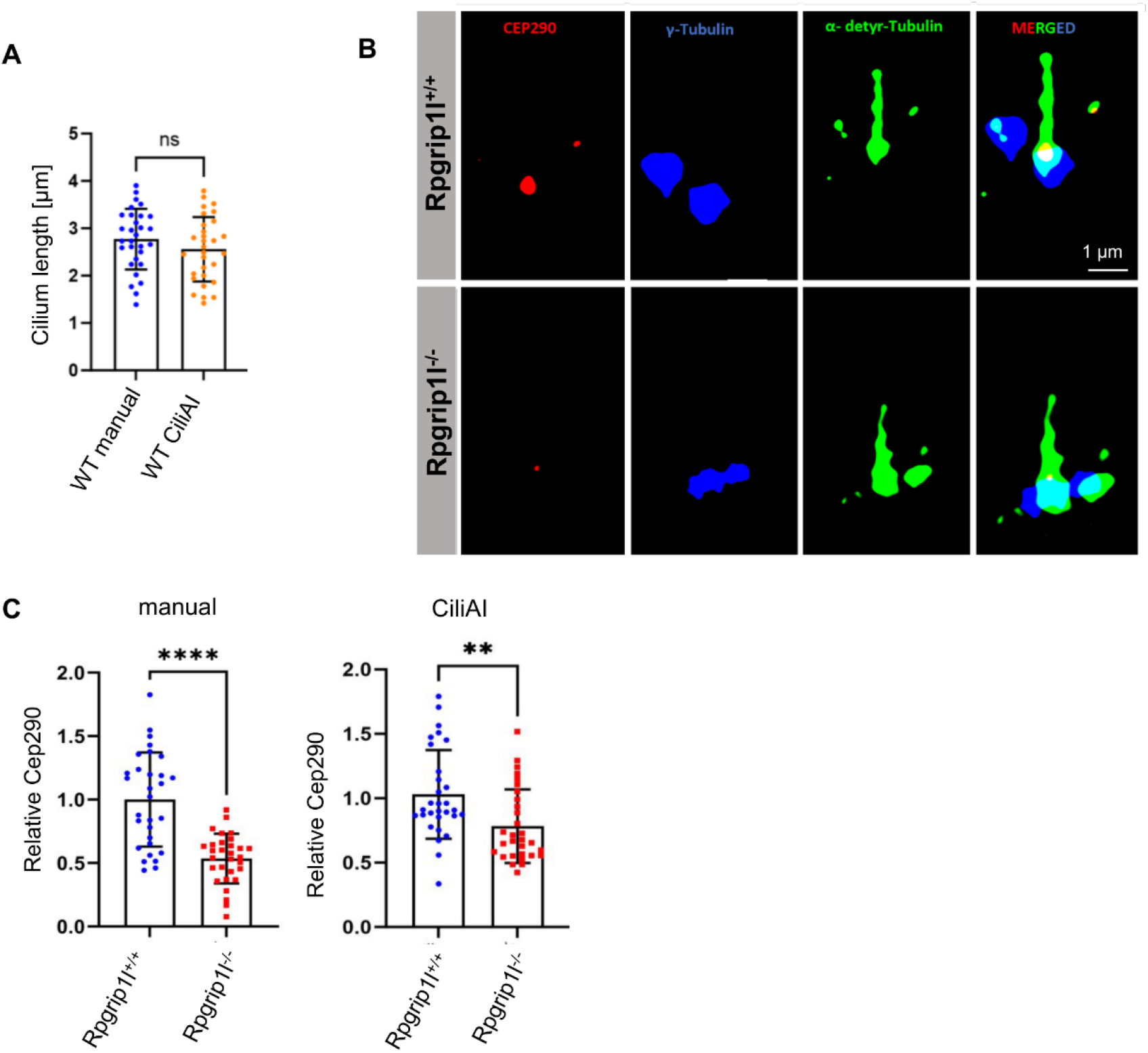
Validation of automated cilium length measurements and densitometry by CiliAI compared with manual analysis. (A) Quantification of cilium length measured manually using Olympus software and automatically using CiliAI. (B) Representative confocal microscopy image of NIH-3T3 *Rpgrip1l*^+/+^ and *Rpgrip1l*^−/−^ cells stained for Cep290 (red, transition zone), γ-tubulin (blue, basal body), and detyrosinated α-tubulin (green, axoneme). Scale bar: 1 µm. (C) Quantification of relative fluorescence intensity within the ciliary transition zone by manual densitometric analysis (left) and CiliAI-based automated analysis (right), showing comparable results between both methods. Data are presented as mean ± SEM. ns = not significant (two-tailed Student’s t-test). Manual = traditional ImageJ-based analysis; CiliAI = AI-based automated quantification.

To evaluate compartment-specific fluorescence quantification, Cep290 localization was analyzed in *Rpgrip1l*^−/−^ and *Rpgrip1l*^+/+^ cells ^22^. *Rpgrip1l*^+/+^ and *Rpgrip1l*^−/−^ cells were immunostained for Cep290, γ-tubulin, and α-detyrosinated tubulin with representative images shown in **Figure 5B**. Manual quantification of Cep290 fluorescence intensity revealed a significant reduction of Cep290 in *Rpgrip1l*^−/−^ cells compared with *Rpgrip1l*^+/+^ controls (**Figure 5C**, left graph). Automated quantification using CiliAI confirmed this reduction (**Figure 5C**, right graph), although the effect size was slightly less pronounced compared with manual evaluation. Automated analysis reproduced previously reported reductions in transition zone-associated Cep290 signal intensity in *Rpgrip1l*-deficient cells ^22^.

To further evaluate automated densitometric analysis in a condition lacking a known ciliary phenotype, Rpgrip1l fluorescence intensity was analyzed in *Rmnd5a*^+/+^ and *Rmnd5a*^−/−^ cells. *Rmnd5a*^+/+^ and *Rmnd5a*^−/−^ cells were immunostained for Rpgrip1l, γ-tubulin, and detyrosinated α-tubulin, with representative images shown in **Figure 6A**. Manual quantification revealed no significant difference in Rpgrip1l fluorescence intensity between genotypes (**Figure 6B**, left graph). Automated analysis using CiliAI similarly detected no significant difference in Rpgrip1l signal intensity between genotypes (**Figure 6B**, right graph). Together, these findings indicate that CiliAI reproduced both positive and negative experimental outcomes observed by manual densitometric analysis.

**Figure 6:**
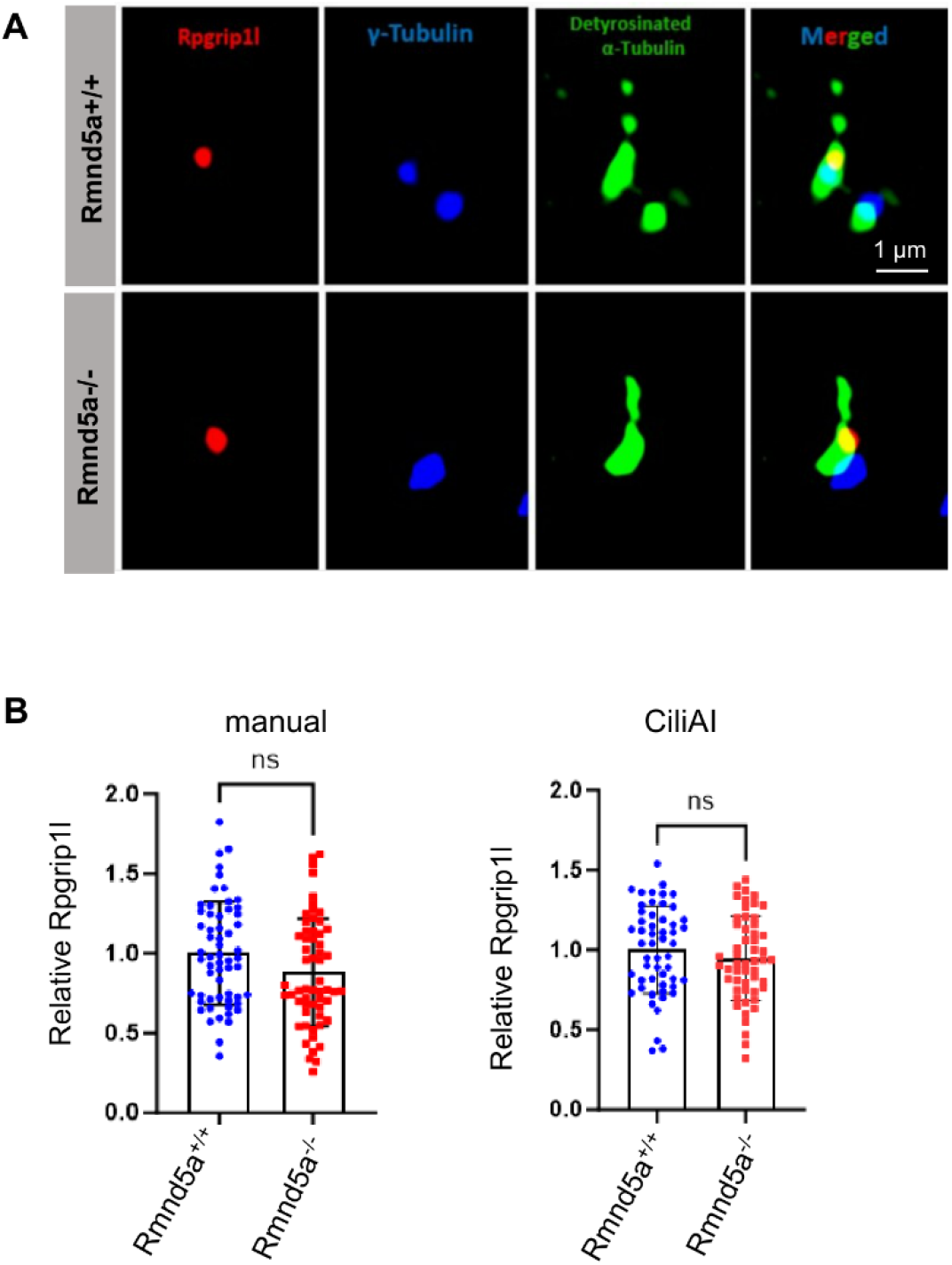
Validation of automated ciliary Rpgrip1l densitometry in *Rmnd5a*^+/+^ and *Rmnd5a*^−/−^ cells. (A) Representative confocal microscopy images of *Rmnd5a*^+/+^ and *Rmnd5a*^−/−^ cells stained for Rpgrip1l (red, transition zone marker), γ-tubulin (blue, basal body marker), and detyrosinated α-tubulin (green, axoneme marker). Merged panels show co-localization of ciliary substructures. Scale bar: 1 µm. (B) Quantification of relative Rpgrip1l fluorescence intensity in *Rmnd5a*^+/+^ and *Rmnd5a*^−/−^ cells by manual analysis (left) and automated CiliAI analysis (right). Both genotypes. Data are presented as mean ± SEM. ns = not significant (two-tailed Student’s t-test). Manual = traditional ImageJ-based densitometric analysis; CiliAI = AI-based automated quantification.

## 3 Discussion

CiliAI was developed to address a central limitation in quantitative cilia analysis: the need for reproducible detection, substructure segmentation, and compartment-specific fluorescence quantification from multiplexed confocal microscopy datasets. Manual analysis of ciliary morphology and protein abundance remains labor-intensive and user-dependent, particularly when measurements are performed across z-stacks and require selection of small subciliary regions. By integrating automated cilia localization, segmentation of the basal body, transition zone, and axoneme, and fluorescence quantification within these compartments, CiliAI provides an integrated workflow for standardized cilia analysis.

The AI models achieved performance sufficient to support automated downstream analysis of primary cilia. For cilia detection, the model reached an mAP50–95 of 67.5% for segmentation masks and 70.0% for bounding boxes, while AP50 exceeded 94%, indicating reliable object localization despite the small size and morphological variability of ciliary structures. Substructure segmentation achieved a mean AP50 of 90.5% and a mean AP50–95 of 58.1% across basal body, transition zone, and axoneme classes. Performance was highest for axoneme and basal body segmentation, whereas transition-zone segmentation remained more challenging because of its small size and variable morphology. These findings indicate that the models provide sufficient spatial precision for automated morphometric and fluorescence-based analyses while identifying transition-zone segmentation as a key area for future model refinement.

Biological validation further showed that CiliAI reproduced manual measurements in independent analytical contexts. Automated cilium length measurements in NIH-3T3 cells showed no statistically significant difference from manual quantification. In addition, CiliAI reproduced the reduction of transition zone-associated Cep290 signal intensity in *Rpgrip1l*-deficient cells, consistent with previous findings ^22,24^. In contrast, both manual and automated analysis detected no significant change in Rpgrip1l fluorescence intensity in *Rmnd5a*-deficient cells. The ability to reproduce both positive and negative experimental outcomes is important, because automated analysis should not only detect known phenotypes but also avoid generating apparent differences in conditions without measurable changes.

CiliAI extends existing cilia-analysis workflows by combining cilia detection, substructure segmentation, and compartment-specific fluorescence quantification in a single web-based pipeline. Existing tools such as CiliaQ ^26^ and ACDC ^27^ have enabled important advances in automated cilia detection and morphometric analysis, and newer workflows appear to support ciliary subdomain analysis ^28^. CiliAI complements these approaches by focusing on automated analysis of ciliary substructures and quantitative fluorescence measurements from multiplexed confocal image data. This may be useful for studies of ciliary protein localization, proteostasis-associated phenotypes, and standardized image-based screening workflows.

Several limitations remain. First, the current implementation is optimized for Olympus .oir confocal files, and broader compatibility with additional microscopy formats will be required for wider adoption. Second, model performance may decrease for image types, staining conditions, cell types, or microscopy platforms underrepresented in the training data. Future development should therefore include multi-laboratory datasets and external validation across different imaging systems. Third, although the current validation demonstrates agreement with manual analysis for selected morphometric and densitometric readouts, broader benchmarking against established cilia-analysis tools and additional biological conditions will be needed to define the generalizability of the workflow. Future studies should include detailed annotation statistics, dataset composition, and segmentation performance across independent test sets to further strengthen reproducibility and support broader deployment.

## 4 Materials and Methods

### 4.1 Cell culture, immunofluorescence, and image acquisition

NIH-3T3 cells (ATCC Cat# CRL-6442, RRID:CVCL_0594) were maintained in Dulbecco’s modified Eagle’s medium (DMEM) supplemented with 10% (v/v) fetal calf serum (FCS) and 4500 mg/l glucose (high concentration) if not mentioned otherwise. For detection of primary cilia, NIH-3T3 cells were grown to confluency and serum-starved with medium containing 0.5% FCS for at least 24 h on glass coverslips (Epredia, P231.1). Cells were fixed with 4% PFA (1 h at 4°C) for Rpgrip1l and Arl13b or with methanol for (5 min at −20°C) for Cep290 specific staining, rinsed three times with PBS, followed by permeabilization with PBS/0.5% Triton X-100 for 10 min. After three washes with PBS/0.1% Triton X-100, cells were blocked at room temperature in PBS/0.1% Triton X-100 containing 10% donkey serum for 1 h. Diluted primary antibodies in PBS/0.1% Triton X-100 1% blocking solution were incubated overnight at 4°C. After three washing steps with PBS/0.1% Triton X-100, incubation with secondary antibody in PBS/0.1% Triton X-100 blocking solution was performed three times at room temperature for 10 min and subsequent mounting with Mowiol (ROTH, #0713.2). Image acquisition of single plane images was carried out using an Olympus confocal microscope (Olympus FV3000) with TruSight (deconvolution) technology with the following settings: DAPI 430-470 nm 5% intensity, Alexa Flour 488 500-540 nm 2% intensity, Alexa Flour 568 570-670 nm 2% intensity. A 100 × oil-immersion objective (Olympus N5702400) with a numerical aperture of 1.45 was used. The polyclonal antibody against Rpgrip1l was generated by immunizing rabbits with a GST-Rpgrip1l fusion protein encompassing the murine Rpgrip1l-RID domain (amino acids 1,060–1,264) by Pineda antibody services.

| Antibody | Source | Order number | Species | Dilution |
| --- | --- | --- | --- | --- |
| Ac-Tubulin | Sigma-Aldrich | T6793 | mouse | 1:100 |
| Ac-Tubulin | Cell Signaling Technology | 5335 | rabbit | 1:100 |
| γ-Tubulin | Sigma-Aldrich | T6557 | mouse | 1:200 |
| γ-Tubulin | Santa Cruz Biotechnology | sc-7396 | mouse | 1:50 |
| Arl13b | Proteintech | 17711-1-AP | rabbit | 1:100 |
| Cep290 | Proteintech | 22490-1 | rabbit | 1:200 |
| Tmem67 | Proteintech | 13975-1-AP | rabbit | 1:100 |
| detyrosinated α-tubulin | EMD Millipore | AB3201 | rabbit | 1:50 |
| Rpgrip1l | Self made <sup>29,30</sup> | - | rabbit | 1:200 |
| Anti-goat Delight488 | Jackson ImmunoResearch | 705-475-003 | donkey | 1:100 |
| Anti-mouse Dylight405 | Jackson ImmunoResearch | 715-475-150 | donkey | 1:50 |
| Anti-rabbit Cy3 | Jackson ImmunoResearch | 711-165-152 | donkey | 1:100 |

### 4.2 Manual quantification

Intensity of ciliary proteins based on immunofluorescence staining was measured as described before by using ImageJ ^21,22,24,31^. For the quantification of Arl13b, we used the entire ciliary membrane area marked by detyrosinated α-tubulin and quantified the average pixel intensity to take cilia length into account thereby receiving the data of WT cilia. For all other ciliary protein intensities, we selected the region labelled by γ-tubulin (for Rpgrip1l) or the area in-between the γ-tubulin staining and the proximal part of the detyrosinated α-tubulin staining (for TZ proteins) and measured the total pixel intensity. The labelling of the selected region was outlined with the freehand selection tool and the mean intensity of the desired channel inside the area measured in an 8-bit scale (0-255). To get rid of the ratio of unspecific (background) staining, we subtracted the mean value of the average pixel intensity (in the case of Arl13b) or of the total pixel intensity (for Rpgrip1l and TZ proteins) of three neighbouring regions free from specific staining. The quantified areas of the specific staining and the unspecific staining of every individual measurement were equal in size.

### 4.3 Statistical analysis

The statistical analysis of the data was performed using GraphPad Prism. Data are presented as mean ± standard error of mean (SEM). Two-tailed Student’s *t-*test was performed for all data in which two datasets were compared. *P*-value < 0.05 was considered to be statistically significant, *P*-value < 0.01 was regarded as statistically very significant, and *P*-value < 0.001 was accounted statistically highly significant.

### 4.4 Training dataset preparation and annotation

For the AI-driven detection and analysis pipeline, 3D confocal image stacks were prepared as described in Section 2.1. Every fifth z-stack was extracted to minimize redundancy, yielding approximately 500 unique 2D images from over 50 original 3D stacks acquired using an Olympus FV3000 confocal microscope. For cilia localization, full-frame 1024 × 1024 px confocal images were subdivided into four overlapping tiles using a 2×2 grid with 20% overlap. This tiling strategy enhances feature learning by providing multiple overlapping views of each cilium and ensuring that cilia located near tile boundaries are captured completely in at least one tile. The annotated tile sets were uploaded to the Datamarkin platform, leveraging the Segment Anything Model (https://arxiv.org/abs/2304.02643) for expert-assisted annotation.

In the second AI model for substructure analysis, individual cilia were cropped from the confocal images. Given their small size (20–60 px in the original images), pre-processing included adding a margin (10% of the longest side) and uniform padding to create square images, which were then resized to 1024 × 1024 px using OpenCV’s INTER_CUBIC interpolation to preserve fine structural details. To aid annotation, a composite image was generated for each cilium, displaying the original RGB image alongside isolated red, green, and blue channels, allowing annotators to more clearly distinguish the basal body (blue channel), transition zone (red channel), and axoneme (green channel).

### 4.5 Neural network architecture and training

For automated cilia detection and segmentation, we used Detectron2’s implementation of Mask R-CNN ^32^. with a ResNeXt-101 (X_101_32 × 8d) backbone and Feature Pyramid Network (FPN). Both the cilia-detection and substructure-segmentation models were initialized from the standard mask_rcnn_X_101_32 × 8d_FPN_3x configuration and trained using Detectron2’s default optimization settings. All models were trained using Detectron2’s implementation of Mask R-CNN with a ResNeXt-101-FPN backbone (mask_rcnn_X_101_32 × 8d_FPN_3x). Training was performed using the default configuration provided by Detectron2 for this architecture, without architecture-specific modifications or custom hyperparameter optimization.

### 4.6 Inference workflow and image processing

Once the user uploads a raw 3D confocal imagery (Olympus OIR format), the CiliAI platform automatically inspects the file using ImageJ to determine the number of z-stacks, image dimensions, and bit depth. In some cases, the original file metadata cannot be read programmatically, requiring manual entry of key parameters such as the color-channel order, pixel size calibration (nm/pixel), and inter-slice spacing (z-distance). Where possible, default values are pre-filled to reduce user input requirements.

#### 4.6.1 Slice extraction and normalization

Each 3D imagery is decomposed into its constituent 2D z-stack slices. To enable consistent analysis across different imaging sessions and bit depths, pixel intensities are normalized. The platform automatically detects the bit depth (8-bit, 12-bit, or 16-bit) by examining the maximum pixel value in the imagery. User-specified channel ordering ensures that red, green, and blue channels correspond correctly to the biological markers used (typically: blue = basal body/γ-tubulin, red = transition zone/RPGRIP1L, green = axoneme/acetylated tubulin).

Background intensity is calculated to enable normalization of fluorescence signals across different imaging sessions and microscope settings. We define background as the mean intensity of the lowest 2% of pixel values across the entire 3D image stack. This background value is subsequently subtracted from all density measurements to provide background-corrected fluorescence intensities.

#### 4.6.2 Tiling and overlap strategy

To improve detection accuracy and avoid missing small cilia, each z-stack slice (1024 × 1024 px) is partitioned into overlapping tiles using a 2×2 grid with 20% overlap, generating 4 tiles per slice. Each tile has dimensions of approximately 614 × 614 px from the original image. This overlap strategy ensures that cilia falling near tile boundaries are captured completely in at least one tile, preventing edge-related detection failures.

After cropping, each tile is upscaled to 1200 × 1200 pixels to increase the effective resolution for the detection network. This upscaling step is particularly important for detecting fine ciliary structures that might occupy only 20–60 pixels in the original image. By focusing the AI model on smaller, upscaled regions, cilia that might otherwise be overlooked in a single full-frame image are more reliably detected. The overlapping regions between tiles ensure that the same cilium detected in multiple tiles can be subsequently merged using IoU-based non-maximum suppression.

#### 4.6.3 Primary cilia Detection

The first AI model processes each of the four tiles independently to identify potential cilia. The model outputs bounding box coordinates, confidence scores (0–1 scale), and segmentation masks for each detected cilium. Since overlapping tiles create redundant detections for cilia appearing near tile boundaries, a merging step is required.

Detected cilia are transformed back to whole-image coordinates using the known tile positions and upscaling factors (614 px original tile → 1200 px processed tile). Duplicate detections are identified and merged using the Intersection-over-Union (IoU) metric (Supplementary Figure 1). For each pair of detections, IoU is calculated as the area of overlap divided by the area of union between their bounding boxes. Detection pairs with IoU ≥ 0.5 are considered to represent the same cilium and are merged by retaining the detection with higher confidence score. This non-maximum suppression (NMS) procedure eliminates duplicates while preserving all unique cilia across the image.

#### 4.6.4 Substructure segmentation

After global localization of cilia, individual detections are extracted for higher-resolution substructure analysis. Each detected cilium is cropped from the original image data with a padding margin of 10% (relative to the longest dimension of the bounding box) to ensure complete capture of the ciliary structure. The cropped region is expanded to a square shape by adding uniform padding, then resized to 1024 × 1024 pixels using OpenCV’s INTER_CUBIC interpolation to preserve structural details.

These high-resolution cilium images are passed to the second AI model that performs instance segmentation of three ciliary substructures: the axoneme (in our case typically marked with anti-acetylated or anti-detyrosinated tubulin antibodies in the green channel), the basal body (typically marked with anti-γ-tubulin antibodies in the blue channel), and the transition zone (typically marked with antibodies against proteins like Rpgrip1l or Cep290 in the red channel). The model outputs detailed segmentation masks delineating each substructure.

To ensure biological validity, an optional validation step verifies that each detected cilium contains at minimum both an axoneme and a basal body, as these are the defining structural features of a primary cilium. Detections lacking either component can be flagged or filtered depending on user preferences.

#### 4.6.5 Quantitative measurements

##### Fluorescence Density Analysis

For each segmented substructure, fluorescence intensity is measured within the corresponding mask region. To enable comparison across different microscopes and imaging conditions, raw intensity values are retrieved directly from the original 3D image data (not from the processed/tiled images) and normalized by the maximum pixel value for the detected bit depth (255 for 8-bit, 4095 for 12-bit, 65535 for 16-bit). The pre-calculated background intensity is then subtracted from the normalized value. This yields a background-corrected, normalized density measurement on a 0–1 scale that designed to facilitate comparison across samples and imaging sessions.

Channel-specific analysis assigns fluorescence measurements to the appropriate biological structure: axoneme intensity is measured in the green channel, basal body intensity in the blue channel, and transition zone intensity in the red channel, corresponding to the typical fluorophore-antibody combinations used in cilia immunofluorescence protocols.

##### Morphometric Measurements

Cilium length was approximated as the maximal internal distance within the axoneme segmentation mask. This approach accommodates curved or bent cilia, as it measures the longest interior path rather than assuming a straight structure. The areas of the basal body, transition zone, and axoneme are calculated from their respective segmentation masks using the Shoelace formula for polygon area. Because these measurements are initially in pixel coordinates, they are converted into metric units (nanometers for length, square nanometers for area) using the pixel size calibration specified by the user or derived from file metadata.

To optimize data storage and transmission, segmentation masks are simplified using the Ramer-Douglas-Peucker algorithm with a tolerance of 1 pixel. This reduces the number of polygon vertices while preserving geometric accuracy, decreasing file size by approximately 70–80% without affecting measurement precision.

#### 4.6.6 Z-stack integration and output generation

CiliAI links 2D detections across z-stack slices to summarize cilia detected across the imaging volume. Cilia appearing in multiple consecutive z-stacks are tracked and linked based on their spatial overlap. For each detected cilium in the summary dataset, we identify all z-stack slices in which a spatially overlapping detection occurs (IoU ≥ 0.5). This generates a “visibility array” indicating the z-stack range across which each cilium extends, providing insight into its three-dimensional extent through the imaging volume.

A 2D merged “top view” projection of the entire volume is generated by applying NMS across all z-stacks with an IoU threshold of 0.5, creating a composite map that highlights the spatial distribution of all detected cilia in the field of view. This projection facilitates rapid quality control and provides an intuitive overview of ciliary organization across the cell population.

### 4.7 Model performance evaluation

Both models were evaluated on a held-out validation set comprising 10% of the annotated data, split at the level of source 3D image stacks so that no tile or cilium from a validation stack appeared in training. The cilia-detection validation set comprised 28 tiles (63 annotated cilia); the substructure-segmentation validation set comprised 32 cilium crops (118 annotated substructures: 39 basal body, 45 transition zone, 34 axoneme). These per-class counts refer to the expert-annotated ground-truth instances in the held-out validation set against which model predictions were compared; all reported metrics were computed on this validation set, and no independent external test set was used. Detection and segmentation performance were quantified using standard COCO metrics: average precision at an IoU threshold of 0.50 (AP50), at 0.75 (AP75), and averaged across IoU thresholds from 0.50 to 0.95 in 0.05 steps (AP). AP50 reflects object-level detection with lenient localization, whereas AP and AP75 require progressively stricter spatial agreement between predicted and annotated masks. The cilia-detection model was evaluated as a single object class (cilium). The substructure model was evaluated per class (basal body, transition zone, axoneme) and reported as both the mean across classes and the individual per-class values. Metrics were computed using the COCOEvaluator implementation in Detectron2.

AP50 is the average precision computed at a single IoU threshold of 0.50, i.e. a predicted instance counts as correct when it overlaps the matching ground-truth annotation by at least 50%. AP75 applies the same definition at the stricter IoU threshold of 0.75. The mean average precision (AP, equivalent to mAP50–95) is the average of per-threshold AP values across ten IoU thresholds from 0.50 to 0.95 in steps of 0.05, and therefore rewards progressively more precise boundary localization. Pixel-level mask accuracy and the per-pixel false-positive and false-negative rates (Figure 4E,F) were taken directly from the Detectron2 training log (mask_rcnn/accuracy, mask_rcnn/false_positive, mask_rcnn/false_negative fields), which compare predicted foreground masks against ground-truth masks for matched instances at each checkpoint.

## 5 Data availability

The raw confocal microscopy datasets (Olympus .oir files) acquired in this study are available from the corresponding authors on reasonable request for non-commercial academic use.

## 6 Code availability

The CiliAI deployment codebase, trained model weights, and Detectron2 training logs underlying the performance metrics reported in Figure 4 are available at https://github.com/datamarkin/ciliai, released free of charge for non-commercial academic use under the accompanying license.

## 7 Acknowledgements

We thank Ilona Renken-Olthoff and the entire team of the Health and Medical University for their support. T.P. was supported by Deutsche Forschungsgemeinschaft under Grant GRK 2155 (ProMoAge) and by DFG grant 566709590.

## 8 Author contributions

TP, CG and EN conceived the study and designed experiments. CG and EK performed immunofluorescence studies with subsequent intensity quantifications, cilia length measurements and statistics. TP, CG and EK prepared figures. TP and EN wrote the manuscript with assistance from CG. All authors approved the manuscript. ChatGPT was used for language editing.

## 9 Competing Interests

E.N. is affiliated with Datamarkin, the company that developed the AutoML platform and the CiliAI web application described in this study. The remaining authors declare no competing interests.

## 10 Figure Legends

**Supplementary Figure 1:**
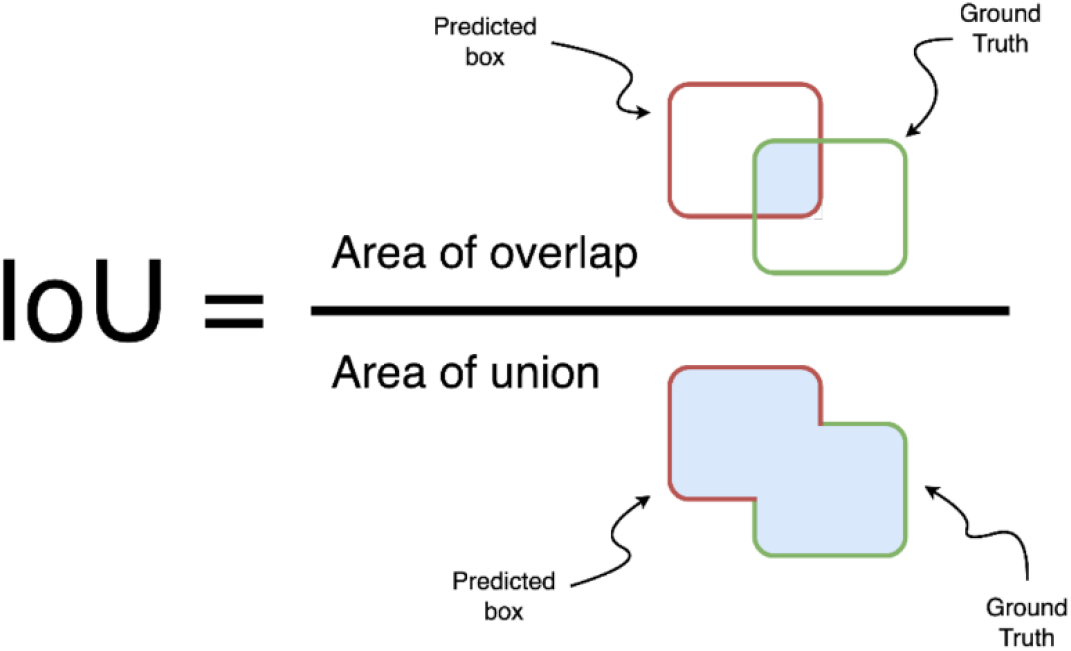
Concept of Intersection over Union (IoU) used for evaluating cilia detection accuracy. Illustration of the Intersection over Union (IoU) metric used in object detection to quantify the overlap between the predicted cilium bounding box (red) and the ground truth annotation (green). IoU is defined as the ratio of the area of overlap between the two boxes to the area of their union. Higher IoU values indicate closer agreement between model predictions and expert annotations.

## Notes

https://github.com/datamarkin/ciliai

